# The Parasite Extinction Assessment & Red List: an open-source, online biodiversity database for neglected symbionts

**DOI:** 10.1101/192351

**Authors:** Colin J. Carlson, Oliver C. Muellerklein, Anna J. Phillips, Kevin R. Burgio, Giovanni Castaldo, Carrie A. Cizauskas, Graeme S. Cumming, Tad A. Dallas, Jorge Doña, Nyeema Harris, Roger Jovani, Zhongqi Miao, Heather Proctor, Hyun Seok Yoon, Wayne M. Getz

**Affiliations:** Department of Environmental Science, Policy, and Management, University of California Berkeley; Berkeley, CA, United States.; Department of Invertebrate Zoology, National Museum of Natural History, Smithsonian Institution; Washington, DC, United States.; Department of Ecology and Evolutionary Biology, University of Connecticut; Storrs, CT, UnitedStates.; ARC Centre of Excellence in Coral Reef Studies, James Cook University;Townsville, Queensland, Australia.; Department of Environmental Science and Policy,University of California; Davis, CA, United States.; Department of Evolutionary Ecology,Estación Biológica de Doñan(CSIC); Americo Vespucio s/n, E-41092 Sevilla, Spain.; Ecology and Evolutionary Biology, University of Michigan; Ann Arbor, MI, United States.; Department of Biological Sciences, University of Alberta; Edmonton, Alberta, Canada.; School of Mathematical Sciences, University of KwaZulu-Natal; South Africa.

**Author notes:** For correspondence (CJC); (OCM). These authors contributed equally to this work (Colin J. Carlson, Oliver C. Muellerklein).

## Abstract

Parasite conservation is a rapidly growing field at the intersection of ecology, epidemiology, parasitology, and public health. The overwhelming diversity of parasitic life on earth, and recent work showing that parasites and other symbionts face severe extinction risk, necessitates infrastructure for parasite conservation assessments. Here, we describe the release of the Parasite Extinction Assessment & Red List (PEARL) version 1.0, an open-access database of conservation assessments and distributional data for almost 500 macroparasitic invertebrates. The current approach to vulnerability assessment is based on range shifts and loss from climate change, and will be expanded as additional data (e.g., host-parasite associations and coextinction risk) is consolidated in PEARL. The web architecture is also open-source, scalable, and extensible, making PEARL a template for more efficient red listing for other high-diversity, data-deficient groups. Future iterations will also include new functionality, including a user-friendly open data pository and automated assessment and re-listing.

## Introduction

Parasitism is one of the most common forms of life on Earth, if not, by species totals, the most common (***Larsen et al., 2017; Dobson et al., 2008***). Even excluding microparasites—such as some bacteria, viruses, and protozoans—the remaining diversity of macroparasites (e.g., helminths, leeches, lice, ticks, fleas, and mites) is staggering and comparatively understudied in ecology. Basic questions remain open at the intersection of parasitology with other fields like community ecology (***Johnson et al., 2015***), evolutionary ecology (***Morand, 2015***), macroecology (***Stephens et al., 2016***), and climate change biology (***Brooks and Hoberg, 2007***; ***Altizer et al., 2013***; ***Cizauskas et al., 2017***). As the field of disease ecology has matured, a growing body of work has shown that parasites are a critical part of ecosystems, acting as regulators of food webs and host populations, and serving an important role in energy 2ow through trophic levels (***Dunne et al., 2013***). The increasingly-apparent bene1ts of parasites make a case for their recognition as an important neglected target for conservation (***Whiteman and Parker, 2005***; ***Pizzi, 2009***; ***Carlson et al., 2013***; ***Gómez and Nichols, 2013***; ***Dougherty et al., 2016***; ***Rocha et al., 2016***), especially given that parasitic life cycles are already known to be particularly extinction-prone due to cascading co-extinctions with hosts (***Durden and Keirans, 1996***; ***Dunn et al., 2009***; ***Dallas and Cornelius, 2015***; ***Farrell et al., 2015***). With recent work showing that climate change and coextinction combined could threaten one in every three helminth parasite species (***Carlson et al., 2017***), frameworks to assess and catalog parasite extinction risk are urgently needed.

While institutions such as the IUCN have spent decades developing centralized frameworks for prioritizing the conservation of free-living biodiversity, parasites are rarely included in main-stream assessments; for example, only two animal macroparasites are listed on the IUCN Red List (*Hematopinus oliveri*, the pygmy hog louse, and *Hirudo medicinalis*, the medicinal leech). The under-representation of parasites speaks to broader deficiencies in IUCN invertebrate assessments, but also to the comparative bias against parasites in conservation, which has conventionally treated parasites and disease as synonymous (***Dougherty et al., 2016***). Mainstreaming parasites into conservation requires researchers to address a number of additional factors: the independent extinction risk of parasites as well as coextinction risk, the degree of host specificity, the modes and efficiency of transmission, the possibility for unintended consequences to wildlife or human health, the cost-effectiveness of parasite conservation as a compatible goal with host conservation, and the feasibility of *ex situ* conservation (***Dougherty et al., 2016***).

At the most basic level, incorporating parasite conservation measures into existing free-living species’ conservation plans can prevent accidental or deliberate loss of affiliates (***Jørgensen, 2015***),the extinction of the California condor’s louse *Colpocephalum californianus*, or the black-footed ferret louse *Neotrichodectes sp.* (***Stringer and Linklater, 2014***). Some key assessments have been made for parasites of high-profile hosts like the black-footed ferret (***Gompper and Williams, 1998***) or the Tasmanian devil (***Wait et al., 2017***), but more expansive assessments are rare. Recent work has pushed to embrace a broader perspective on symbiosis within parasitology (***Jovani et al., 2017***), and in the context of global change biology, we believe this is an important step towards effective conservation. Symbionts as a broad group face a common set of challenges, and the same conservation measures that make sense for parasites are part of a broader shift towards symbiont-conscious conservation.

Rising interest in parasite conservation comes at a time when open data in parasitology is rapidly expanding; while host-parasite association data have historically been available from scientific collections or online databases (***Strona and Lafferty, 2012***; ***Dallas, 2016***), only in the last few years have major sources such as the U.S. National Parasite Collection (***Lichtenfels et al., 1992***; ***Carlson et al., 2017***) or Global Mammal Parasite Database (***Stephens et al., 2017***) been updated to include the detailed spatial data that are critical for conservation assessments. Expanding existing repositories, and improving access to collections data, is already a key part of ongoing work bridging the gap between parasitology and other fields like epidemiology, disease ecology, and conservation (***DiEuliis et al., 2016***). For parasite conservation to be operational and actionable in the shortest term, researchers currently need a detailed and dedicated bioinformatic repository for the explicit purpose of centralizing data on population trends, extinction risk, distributions, and conservation efforts. Here, we present the Parasite Extinction Assessment & Red List (PEARL: pearl.berkeley.edu), the first standalone global parasite conservation assessment, database, and web interface.

## PEARL version 1.0

The Parasite Extinction Assessment & Red List compiles the preliminary work of the Parasite Extinction Research Project (2013-2016). The core study (***Carlson et al., 2017***) accomplished three tasks:

1. The georeferencing of the U.S. National Parasite Collection (USNPC), and the compilation of the most detailed occurrence dataset for macroparasites currently available.
2. The estimation of global parasite extinction risk: an estimated 5-10% of species face direct extinction risk from climate change, while up to 30% of helminths face a combined threat from coextinction and climate change.
3. The preliminary release of PEARL version 0.1, a prototype with a static interface, PDF maps, and no search functionality.

Building on PEARL v0.1, the full PEARL v1.0 was released on August 11, 2017 and features an open-source web architecture, which allows a more flexible interface and tool to access the underlying open-access database. Here we formalize the documentation for PEARL v1.0, explaining the updated mechanics of PEARL as a new, standalone form of red listing for parasites and other symbionts; and to outline the use of the framework in the inclusion of parasites in conservation research, and more generally for invertebrate conservation.

## The Assessment

As a scientific resource, PEARL serves two main purposes: an *extinction assessment*, which compiles the extinction risk of enough species that global parasite extinction rates from climate change can be measured; and a *red list*, analogous to the IUCN Red List but focused specifically on measuring the vulnerability of macroparasitic species. The species currently included in the assessment fall into eight major groups: helminth endoparasites (acanthocephalans, cestodes, nematodes, and trematodes) and arthropod ectoparasites (fleas, lice, mites, and ticks). The term parasites is used broadly both here and in the underlying premise of PEARL. While the focus of the assessment is parasitic species, in the overarching goal of mainstreaming parasites into conservation (***Dougherty et al., 2016***), several species and groups are included that are not strictly parasitic. For example, vane-dwelling feather mites (Acariformes: Analgoidea, Pterolichoidea), like many other symbionts, may contextually change roles between parasitism and commensalism or mutualism (***Galván et al., 2012***); within other significant groups in our study, parasitism has secondarily evolved multiple times (***Dorris et al., 1999***; ***Bochkov and Mironov, 2013***).

The vulnerability of species is measured based on projected range loss in the face of climate change, which was forecasted through the use of ecological niche modeling (***Carlson et al., 2017***). Eighteen climate change scenarios are included in that assessment for 457 non-marine macroparasitic species, and extinction risk is estimated based on those projected rates of habitat loss. These are translated into “red listing categories” based on estimated percent range loss in the next 50 years (i.e., by 2070):

- **Critically endangered (CR):** projected decline by ≥ 80% in 50 years
- **Endangered (EN):** projected 50–79% decline in 50 years
- **Vulnerable (VU):** projected 25-49% decline in 50 years
- **Least Concern (LC):** < 25% decline in 50 years

The assessment is designed to be fully transparent about accuracy and bias; accuracy metrics for niche modeling (the true skill statistic and the area under the receiver-operator curve), as well as categorical measures of data quality, are presented alongside distribution models. We developed two data quality metrics based on percentiles within our dataset. Data *coverage* is based on sample size:

| Coverage | Percentile | Occurrence Points |
| --- | --- | --- |
| "Poor" | 0-25 | 0-28 |
| "Fair" | 25-50 | 29-42 |
| "Good" | 50-75 | 43-80 |
| "Excellent" | 75-100 | 81-3289 |

Data *uncertainty* is based on the mean uncertainty radius of every point from manual georeferencing; some species only had GPS-identified data, which were classified as zero uncertainty from georeferencing:

| Uncertainty | Percent | Distance (km) |
| --- | --- | --- |
| "Perfect" | – | 0 |
| "Excellent" | 0-25 | 0-5.1 |
| "Good" | 25-50 | 5.1-6.5 |
| "Fair" | 50-75 | 6.5-7.9 |
| "Weak" | 75-100 | 7.9-21.3 |

By presenting quality as a mainstreamed part of the assessment, we improve transparency, better inform practitioners relying on these data for any applied research, and clearly identify species for which improved data collection is needed (i.e., more precise locality data).

## The Web Interface

PEARL is an open-source web app that builds on several frontend and backend APIs and novel software libraries in both the *Python* and *JavaScript* programming languages. In the backend, PEARL builds upon a **Python 3.5** based web framework developed by *Z. Miao and O. Muellerklein* called Extensible Web App Interactive Mapping, **EWAIM on Github**^1^. EWAIM incorporates continuous unit testing, a basic yet extensible library for server-side *(i.e. backend)* data analysis, and native integration with spatial data structures through **PostGIS / PostgreSQL** or **SQLite**. PEARL’s use of the underlying EWAIM server-side web framework allows near endless GIS functionality to be used on spatial or time series data.

In the frontend, PEARL uses standard *GET / POST* events to process user events to and from the server-side application, allowing interactivity with the backend database via species maps and other functionality in the various web pages / interfaces. Building upon EWAIM, PEARL handles user events to and from web pages through *Flask* protocol, a Python based extensible web microframework. Within the frontend components of PEARL a range of mapping APIs are called. Frontend libraries and API used in PEARL include:

- **D3.js - Data Driven Documents:**^2^ an open-source JavaScript library for dynamic, interactive data visualizations
- **Lealet API:**^3^ an open-source JavaScript library that allows interactive mapping of PEARL tilemaps and provides a mobile-friendly design
- **CARTO Maps API:**^4^ an open-source engine that is scalable and extensible, powers a range of basemaps, and interacts with Leaflet, Google Earth Engine, and CARTO SQL data structures

**Figure 1.**
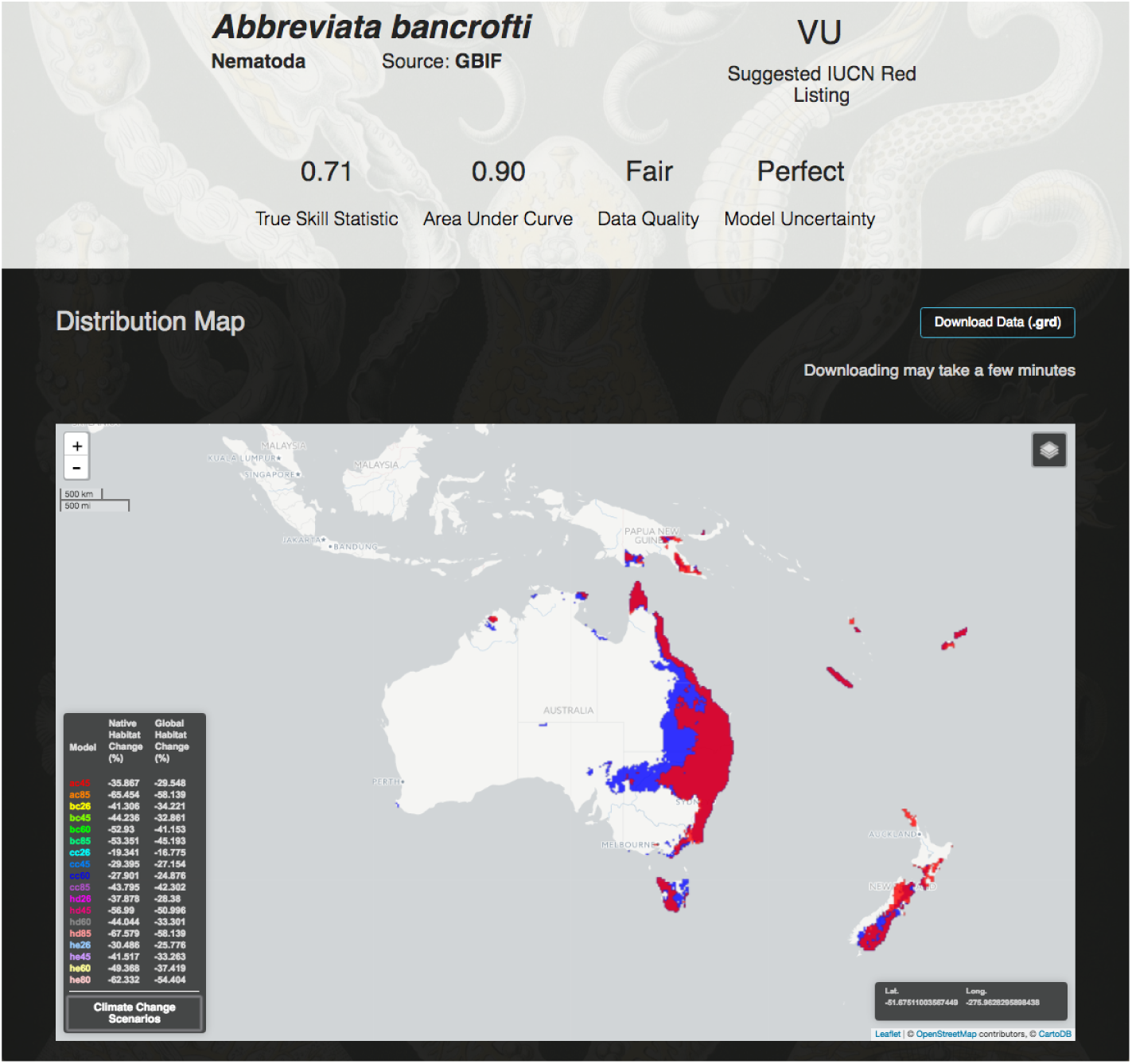
The online interface to PEARL v1.0, illustrated for *Abbreviata bancrofti* (Physalopteridae; Irwin-Smith, 1922), a nematode parasite of the Australian leaf-tailed gecko, *Phyllurus platurus* (Reptilia: Gekkonidae). Results include the current distribution of the nematode (blue) and 18 future climate scenarios. Above, information about the entry is available, including data sources, and model accuracy metrics.

PEARL, as a web app and open-source software, is fully documented and available publicly as a Github repository^5^ with version releases and community-based issues tracked accordingly. The underlying raster files can be downloaded in grid file (.grd) format directly from species pages, and R- and Python-based APIs are currently in development that will allow users to pull data from multiple species at once. Future iterations of PEARL development include, but are not limited to, a more robust mobile-friendly structure; public data uploads and downloads through user login and associated profiles; and the incorporation of a novel dynamic, real-time algorithm that would automate generating and rendering distribution models and associated data analysis based on user data contributions (see Dynamic Updating section below).

## Extending the Framework

The purpose of PEARL is to create a stable platform for parasite conservation that allows continuous improvements by existing teams, and that can incorporate future collaborations from other researchers both in PEARL development and database-building. The full expanded platform makes the interface, and underlying database, extensible in a number of important ways that will be useful both for the future of PEARL, and as a template for the broader problem of invertebrate conservation assessments. After the release of version 1.0, a number of scheduled advances are planned for PEARL over the coming years. We detail five here:

### Assessment II

A major goal for PEARL is the expansion of assessments beyond the 457 species in the pilot study by ***Carlson et al. (2017)***, to include some of the other increasingly-available open data sources in parasitology. The goal of fully georeferencing the U.S. National Parasite Collection–including species with insufficient unique locality points for the main extinction study–by 2018 gives a clear rationale for an expanded assessment, especially in conjunction with the recent release of the Global Mammal Parasite Database v2.0 with spatial data (***Stephens et al., 2017***). Also critical is including smaller, regional datasets from biodiversity hotspots like the Amazon, or the Cape Floristic Region of South Africa. Given that distribution modeling requires a minimum of at least 20-50 occurrences, it would be useful to include species that are currently impossible to map in future iterations by presenting raw occurrence data, rather than niche models. Providing these data may help guide targeted field collection programs that address these data defciencies.

### Reconsidered Criteria

Red listing parasites—or, in fact, any dependent species—poses a more severe methodological problem than already-challenging work on free-living species. In particular, well-designed criteria must accommodate the tremendous diversity of symbiotic groups, and rescale important metrics of viability to an appropriate level; but effective criteria must also (presumably) include information about the vulnerability of hosts. The criteria underlying assessment version 1.0 only indirectly addressed the first of these two challenges, by presenting a radically simplified version of the ***Thomas et al. (2004)*** criteria, which were already reduced from the IUCN Red List criteria.

For comparison, the criteria used in ***Thomas et al. (2004)*** are designed to correspond directly to projected extinction risk:

- **Extinct (EX):** species with a projected future area of zero (100% of species assumed to be committed to eventual extinction)
- **Critically endangered (CR):** projected future distribution area < 10 km^2^, or decline by ≥ 80% in 50 years (species assigned a 75% chance of extinction)
- **Endangered (EN):** projected area 10–500 km^2^, or 50–79% decline in 50 years (species assigned a 35% chance of extinction)
- **Vulnerable (VU):** projected area 501–2,000 km^2^, or > 50% decline in 100 years on the basis of linear extrapolation of 50-year projection (species assigned a 15% chance of extinction)
- **Not Threatened (NT):** 0% extinction risk, no area loss

In the ***Carlson et al.*** (***2017***) study and PEARL v0.1 (and v1.0), these criteria are reduced to percentage-based criteria only:

- **Critically endangered (CR):** projected decline by≥80% in 50 years
- **Endangered (EN):** projected 50–79% decline in 50 years
- **Vulnerable (VU):** projected 25-49% decline in 50 years
- **Least Concern (LC):** < 25% decline in 50 years

The inadequacy of these criteria is a key point articulated by ***Carlson et al.*** (***2017***), and we return to it here to note that it offers only the coarsest level of possible resolution for categorizing extinction risk. More detailed criteria are needed that incorporate risk factors like small ranges (with minimal projected declines), but parasite conservation has yet to develop meaningful benchmarks for these criteria; what is a “small range” for a parasitic species? Is a small range for a trophically-transmitted nematode the same size as a small range for a tick with a single common host? The role of microclimate and heterogeneity within ranges, or of habitat selection and dispersal patterns of hosts, further complicates this problem. Similarly, ***Dougherty et al.*** (***2016***) highlighted the need for advances in population viability analysis for parasites, such that concepts like “minimum viable population” can be readily applied, and included in these criteria.

### Better Host & Parasite Bioinformatics

Host and symbiont extinction risk are fundamentally linked, and conservation of parasites and other symbionts cannot exist in the absence of detailed host information. For ticks and feather mites, underlying data from the ***Carlson et al.*** (***2017***) study contains host association data that can be mainstreamed into future versions. Moreover, access to portals like the helminthR package in R (***Dallas, 2016***) will make it possible to compile detailed information on host-parasite associations for helminths, but these data lack information about the life stage at which different hosts are relevant. Adding life stage-structured data to host-parasite associations will be a key part of PEARL expansion, especially given that host-range disjunctions might be a substantial pressure on parasites in a changing climate (***Pickles et al., 2013***; ***Cizauskas et al., 2017***). A new database published this year makes significant strides towards aggregating life cycle data for acanthocephalans, nematodes, and cestodes (***Benesh et al., 2017***); compiling that data for every species in our study will still likely require the concerted effort of researchers contributing to the expansion of the PEARL database.

The integration of host-parasite association data is an especially sensitive matter in the design of actual parasite conservation schemes. The potential for conflict between parasite conservation and the broader goals of wildlife and human health has already been noted by ***Dougherty et al.*** (***2016***), and a fundamental tenet of effective parasite conservation is attention to potential unanticipated consequences for conservation or public health. Parasites with zoonotic potential or that act as vectors of zoonoses are an especially difficult case, as they may be the target of eradication campaigns simultaneous to conservation efforts for closely related parasites. Developing an aggregated bioinformatic infrastructure that serves both purposes will support parasite research in diverse realms, and facilitate the work of public health practicioners and conservation managers alike. Consequently, future assessments should not only include wildlife (and domestic) host associations, but also detailed information on the known zoonotic potential of every species.

### Integrating Genomic Data

The increasing availability of genomic data associated with the continuous improvements in sequencing and bioinformatics (***Stephens et al., 2015***), and with global initiatives such as the Earth BioGenome project (EBP; ***Pennisi (2017)***), is becoming a huge source of data for conservation assessments (***Pauls et al., 2013***; ***Ikeda et al., 2017***; ***Razgour et al., 2017***). Genomic data can be programmatically gathered from massive databases, such as the NCBI Genome database^6^ and the European Nucleotide Archive (ENA^7^). Notwithstanding, given the bias against symbiont genomes (***Del Campo et al., 2014***), PEARL may need dedicated projects to generate genomic data for the species already included. Doing so open the doors to a number of important new analysies, such as genetically informed ecological niche models (ENMs; ***Marcer et al. (2016)***; ***Ikeda et al. (2017)***) following integrative frameworks (***Razgour et al., 2017***). In this way, assessments can include measures based on neutral and adaptive genomic information to assess the sensitivity of species to environmental variables associated with global change. For these computationally challenging purposes, PEARL will likely require of an increase in computational resources (***Hayden, 2015***).

### Dynamic Updating

One strength of PEARL’s dynamic interface is the potential for continuously-updating red listing, which updates existing species assessments and adds new ones in real-time as new data are contributed by researchers around the world. In an upcoming release, we hope to include an automated tool for continuous integration of new data and assessments, in which submitted spatial data automatically augments the existing global database. This workflow will pave the way for future conservation approaches by allowing dynamically updated red listing, via continuous integration of data into self-updating niche models and respective quality metrics. PEARL can serve as a launching point for an alternative red listing protocol that incorporates machine learning methods to evaluate conservation status at the community or global level (e.g. see recent work by ***Darrah et al. (2017)***), rather than on a manual species-to-species basis (like most current efforts operate), something that will likely be helpful in red listing the 300,000 estimated species of helminths alone.

## Application to Other Conservation Efforts

PEARL is designed to be a template for more successful rapid assessment of conservation risk and status for invertebrates and other difficult-to-profile groups, especially other types of symbionts and coextinction-prone affiliate species. All code for PEARL is publicly available, allowing the rapid and easy development of parallel tools for non-parasitic groups, and encouraging a broader culture of open, reproducible science in conservation. The Github repository contains a detailed user-guide to installing local and server-side instances of PEARL and its underlying web architecture across a number of operating systems, making it easily adaptable for a diverse range of ecological projects. Developing better, broader frameworks for invertebrate conservation could be substantially accelerated with readily available, open-source frameworks for red listing that allow more decentralized data collection and assessment. Data deficiencies for invertebrates are overwhelming (***Clausnitzer et al., 2009***; ***Régnier et al., 2015***), and despite the priority put on red listing insects, there is concern that hyper-diverse groups like the insects will never be described thoroughly enough that conservation assessments can keep pace with extinction rates (***Warren et al., 2007***). Decentralizing the red listing process, and enabling smaller assessments as data are collected, is an important step to protecting not just parasites, but all symbionts and other neglected or understudied groups.

## Acknowledgments

We would like to thank the generous contributions of time and effort to the Parasite Extinction Research Project by our broader team, including Christopher Clements, Eric Dougherty, Sarah Fourby, Nicole Kula, and Sergey Mironov. We especially also thank Gary Casterline for invaluable server and file system assistance for PEARL.

## Footnotes

1 EWAIM on Github: https://github.com/Thru-Echoes/ewaim-webapp

2 D3.js homepage: https://d3js.org/

3 Lea2et API homepage: http://lea2etjs.com/

4 CARTO homepage: https://carto.com/

5 PEARL Github repository: https://github.com/Thru-Echoes/PEARL1.0

6 http://www.ncbi.nlm.nih.gov/genome/

7 http://www.ebi.ac.uk/ena

